# A multi-center study on factors influencing the reproducibility of *in vitro* drug-response studies

**DOI:** 10.1101/213553

**Authors:** Mario Niepel, Marc Hafner, Caitlin E. Mills, Kartik Subramanian, Elizabeth H. Williams, Mirra Chung, Benjamin Gaudio, Anne Marie Barrette, Alan D. Stern, Bin Hu, James E. Korkola, LINCS Consortium, Joe W. Gray, Marc R. Birtwistle, Laura M. Heiser, Peter K. Sorger

**Author notes:** These authors contributed equally (Mario Niepel, Marc Hafner, Caitlin E. Mills). Current address: Ribon Therapeutics, Inc., 99 Hayden Avenue, Building D, Suite 100, Lexington, MA 02421. Current address: Department of Bioinformatics & Computational Biology, Genentech, Inc., South San Francisco, CA 94080. Current address: Clemson University Dept. of Chemical and Biomolecular Engineering Earle Hall 206 S. Palmetto Blvd. Clemson, SC 29634.

## Abstract

Evidence that some influential biomedical results cannot be repeated has increased interest in practices that generate data meeting findable, accessible, interoperable and reproducible (FAIR) standards. Multiple papers have identified examples of irreproducibility, but practical steps for increasing reproducibility have not been widely studied. Here, seven research centers in the NIH LINCS Program Consortium investigate the reproducibility of a prototypical perturbational assay: quantifying the responsiveness of cultured cells to anti-cancer drugs. Such assays are important for drug development, studying cell biology, and patient stratification. While many experimental and computational factors have an impact on intra- and inter-center reproducibility, the factors most difficult to identify and correct are those with a strong dependency on biological context. These factors often vary in magnitude with the drug being analyzed and with growth conditions. We provide ways of identifying such context-sensitive factors, thereby advancing the conceptual and practical basis for greater experimental reproducibility.

## INTRODUCTION

Making biomedical data more findable, accessible, interoperable, and reusable (the FAIR principles (Wilkinson et al., 2016) promises to improve how laboratory experiments are performed and interpreted. Adoption of FAIR approaches also responds to concerns from industrial and academic groups questioning the reproducibility and fundamental utility of biomedical research data (Arrowsmith, 2011; Baker, 2016; Begley and Ellis, 2012; Prinz et al., 2011) and the adequacy of data reporting (Errington et al., 2014; Morrison, 2014). A number of efforts have been launched to repeat published work (https://f1000research.com/channels/PRR), most prominently the Science Exchange Reproducibility Initiative (http://validation.scienceexchange.com/#/reproducibility-initiative). The results of such reproducibility experiments have themselves been controversial (eLIFE-Editorial, 2017; Ioannidis, 2017; Nature-Editorial, 2017; Nosek and Errington, 2017).

In work made possible by the NIH Library of Network-Based Cellular Signatures Program (LINCS) (http://www.lincsproject.org/), this paper investigates the reproducibility of a prototypical class of cell-based experiments rather than focus on a specific published result. This is consistent with the overall goal of LINCS: generating datasets that mesure the responses of cells to perturbation by small molecule drugs, components of the microenvironment, and gene depletion or overexpression. For such a resource to be broadly useful, it must be reproducible. The experiment analyzed in this paper involves determining how tissue culture cells respond to small molecule anti-cancer drugs across a dose range. Such experiments require selection of cell types, assay formats, and time-frames for comparison of pre-and post-treatment cell states; they are therefore prototypical of perturbational biological experiments in general. Drug-response assays are also widely used in preclinical pharmacology (Cravatt and Gottesfeld, 2010; Schenone et al., 2013) and the study of cellular pathways (Barretina et al., 2012; Garnett et al., 2012; Heiser et al., 2012).

In the case of the anti-cancer drugs studied here, cells are typically exposed to drugs or drug-like compounds for several days (commonly three days) and the number of viable cells is then determined, either by direct counting using a microscope or by performing a surrogate assay such as CellTiter-Glo® (Promega), which measures ATP levels in a cell lysate. With some important caveats, the amount of ATP in a lysate from a cell culture dish or well is proportional to the number of viable cells (Tolliday, 2010). Several large-scale datasets describing the responses of hundreds of cell lines to libraries of anti-cancer drugs have recently been published (Barretina et al., 2012; Garnett et al., 2012; Haverty et al., 2016; Seashore-Ludlow et al., 2015), but their reproducibility and utility is being debated (Bouhaddou et al., 2016; CCLE Consortium and GDSC Consortium, 2015; Haibe-Kains et al., 2013; Safikhani et al., 2016).

Our approach was intentionally simple: five experimentally-focused LINCS Data and Signature Generation Centers (DSGCs) would measure the sensitivity of the widely-used, non-transformed MCF10A mammary epithelial cell line to eight small molecule drugs having different protein targets and mechanisms of action. The LINCS Data Coordination and Integration Center (DCIC) would then help to gather the data. One DSGC (hereafter “LINCS Center One”) was charged with studying the possible sources of irreproducibility arising from the inter-Center comparison. Investigators in LINCS Center One had previously shown that conventional drug response measures such as IC_50_ are confounded by variability in rates of cell proliferation arising from variation in plating density, fluctuation in media composition, and intrinsic differences in cell division times (Hafner et al., 2016, 2017a). We corrected for these and other known confounders using the growth rate inhibition (GR) method (Hafner et al., 2016, 2017b; Niepel et al., 2017), thereby focusing the current study on sources of irreproducibility that remain poorly understood. Individual LINCS Centers were provided with identical aliquots of MCF 10A cells, drugs, and media supplements, as well as a detailed experimental protocol and data analysis procedures (Figure 1A). Some variation in the implementation of the protocol was inevitable because not all laboratories had access to the same instrumentation or the same level of technical expertise; in our view, this is a positive feature of the study because it more fully replicates “real-world” conditions.

**Figure 1:**
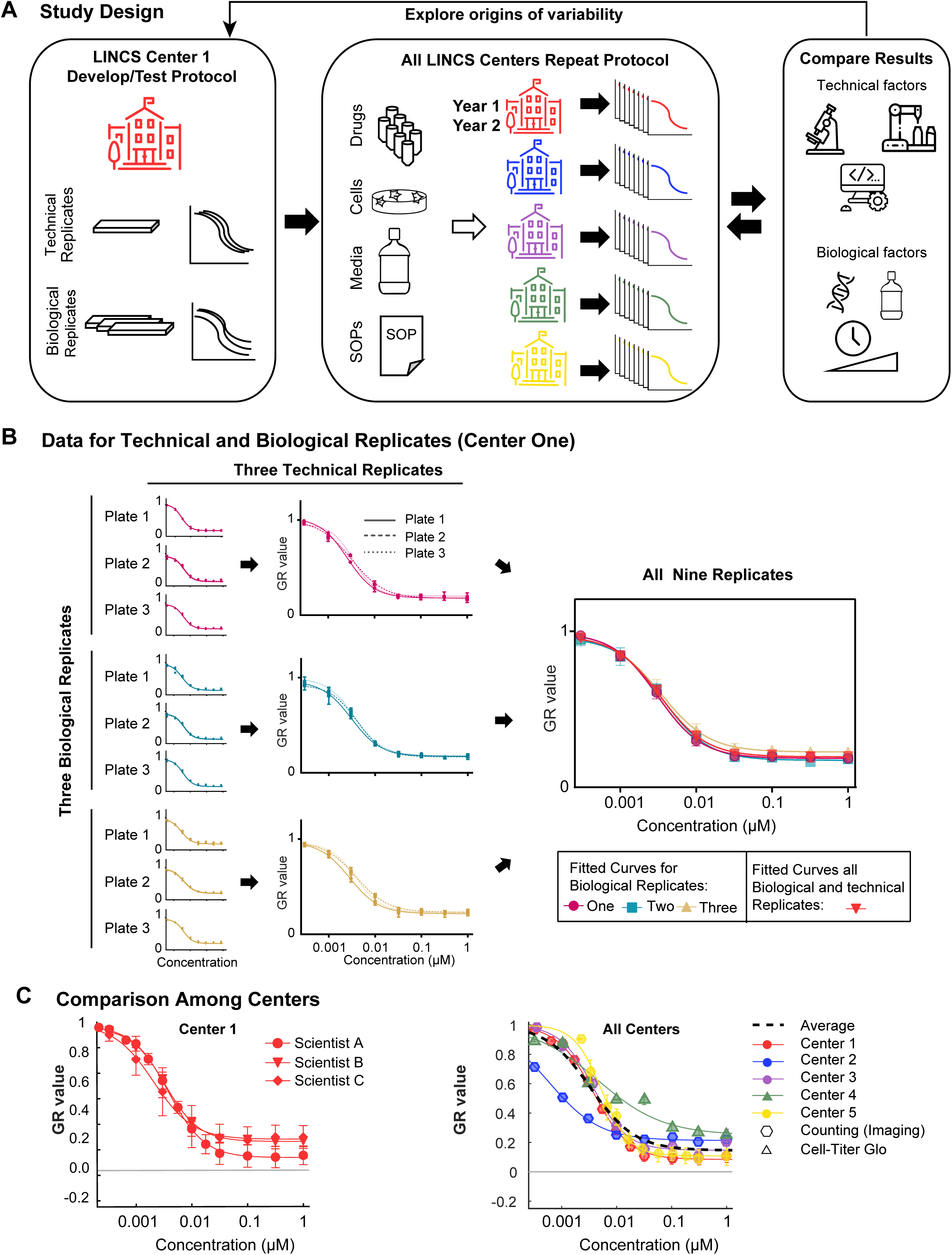
Overview of workflow. (A) LINCS Center One defined the experimental protocol and established within-Center reproducibility by assessment of technical (different wells, plates, same day) and biological (different days) replicates. Common stocks of drugs, cells, and media, as well as a standard experimental protocol was distributed to each of the five data generation centers. Each center performed 72 h dose-response measurements for each of the 8 drugs. LINCS Center One explored the various technical and biological drivers of variability. Technical drivers of variability include assay read-out, use of automation, and analytical pipelines. Biological drivers of variability include cell line isolate, reagents, assay duration, and dose range. This information was fed back to the other Centers to refine their dose-response measurements. (B) Dose-response curves of MCF 10A treated with the MEK1/2 inhibitor Trametinib from a typical experiment showing technical and biological replicates. Technical replicates at the well (triplicate wells per plate), and plate (triplicate plates per experiment) levels make up biological replicates (repeats collected on different days in the same laboratory). The red triangles represent the average of the three biological replicates shown. (C) Independent experiments performed in Center One, and in all Centers (averages of two or more biological replicates). Circles represent the original dataset, triangles represent data collected by a new technician two years after the initial data collection (data shown in B), and diamonds represent data collected as part of a separate LINCS project in Center One http://lincs.hms.harvard.edu/db/datasets/20343/). Inter-Center replicates (averages of one or more biological replicates) performed independently at each LINCS Center. Error bars represent the standard deviation of the mean.

In initial experiments, we observed center-to-center variation in GR50 measurements of up to 200-fold. Center One then performed systematic studies to identify those factors with the largest impact on the measurement of drug response and distributed this information to other centers to improve experimental and analytical procedures. In contrast to several recent studies emphasizing the importance of genetic instability in the variability of cellular phenotypes (including sensitivity to anti-cancer drugs (Ben-David et al., 2018)) we did not find genetic drift to play a significant role in our studies. Instead, irreproducibility arose from a subtle interplay between experimental methods and poorly characterized sources of biological variation as well as differences in data analysis. Based on these findings, we demonstrated that technical staff without previous exposure to our protocol and trained two years after the start of the study could obtain results indistinguishable from assays performed two years previously. Thus, a sustained commitment to characterizing and controlling for variability in perturbation experiments is both necessary and sufficient to obtain reproducible data.

## RESULTS

### Measuring drug responses in collaboration

To establish the single-center precision of dose-response assays, LINCS Center One performed technical and biological replicate measurements using MCF 10A cells and the MEK1/2 kinase inhibitor Trametinib at eight concentrations between 0.33 nM and 1 µM (Figure 1B, C). For technical replicates, multiple drug dilution series were assayed on one or more microtiter plates on the same day. For biological replicates, three sets of assays were performed separated by a minimum of one cell passage in different plates; each biological replicate involved three technical replicates. In all cases, viable cell number was determined by differentially staining live and dead cells, collecting fluorescence images from each well, segmenting images using software, and then counting all viable cells in all wells (Hafner et al., 2016; Niepel et al., 2017). Sigmoidal curves were fitted to the data and four response metrics derived: potency (GR_50_), maximal efficacy (GR_max_), slope of the dose response curve (Hill Coefficient or hGR), and the integrated area over this curve (GRAOC) (Hafner et al., 2016). Fitting procedures and response metrics have been described in detail previously (Hafner et al., 2016, 2017b) (Figure S1A), and all routines and data can be accessed on-line or via download at http://www.grcalculator.org/.

We found that response curves for technical replicates were very similar (Figure 1B), showing that purely procedural error resulting from inaccurate pipetting, non-uniform plating, errors in cell counting etc. were small. Variability in biological replicates as measured by drug potency (log_10_(GR_50_) values) and efficacy (GR_max_ values) was within 1.4 standard deviations for Center One (Figure S2) across three different laboratory scientists.

To measure reproducibility across laboratories while controlling for variation in reagent and genotype, a single LINCS Center distributed to all other Centers identical MCF 10A aliquots, drug stocks, and media additives, as well as a detailed experimental protocol optimized for the cell line-drug pairs under study. This protocol included optimal plating densities, dose-ranges and separation between doses for reliable curve fitting. When individual LINCS centers first performed these assays, up to 200- fold variability in GR50 values was observed (Figure S3). We were surprised by the magnitude of this difference but it is similar to what has previously been observed for large-scale dose-response studies performed by different research teams (Haibe-Kains et al., 2013). To understand the origins of the observed irreproducibility we performed directed and controlled experiments in LINCS Center One.

### Technical drivers of variability

First, we studied the origins of the large inter-Center variability in estimation of *GR_max_* for the topoisomerase inhibitor Etoposide and CDK4/6 inhibitor Palbociclib. We ascertained that one LINCS Center had used the CellTiter-Glo® ATP-based assay and a luminescence plate reader as a proxy for counting the number of viable cells. CellTiter-Glo is among the most commonly used assays for measuring cell viability and it was therefore logical to substitute it for direct cell counting. However, when we performed side-by-side experiments we found that dose-response curves and GR metrics computed from image-based direct cell counts and CellTiter-Glo® were not the same: GR_max_ values for the topoisomerase inhibitor Etoposide and CDK4/6 inhibitor Palbociclib varied by 0.61 and 0.57 respectively for the two assays (GR50 values could not be determined for CellTiter-Glo® data because GR > 0.5 under all conditions tested; Figure 2A). In contrast, in the case of the EGFR inhibitor Neratinib and the PI3K inhibitor Alpelisib, the differences were smaller, varying by 0.03, and 0.24 respectively. This finding likely explains some of the inter-Center differences observed in drug response metrics (Figure S3).

**Figure 2:**
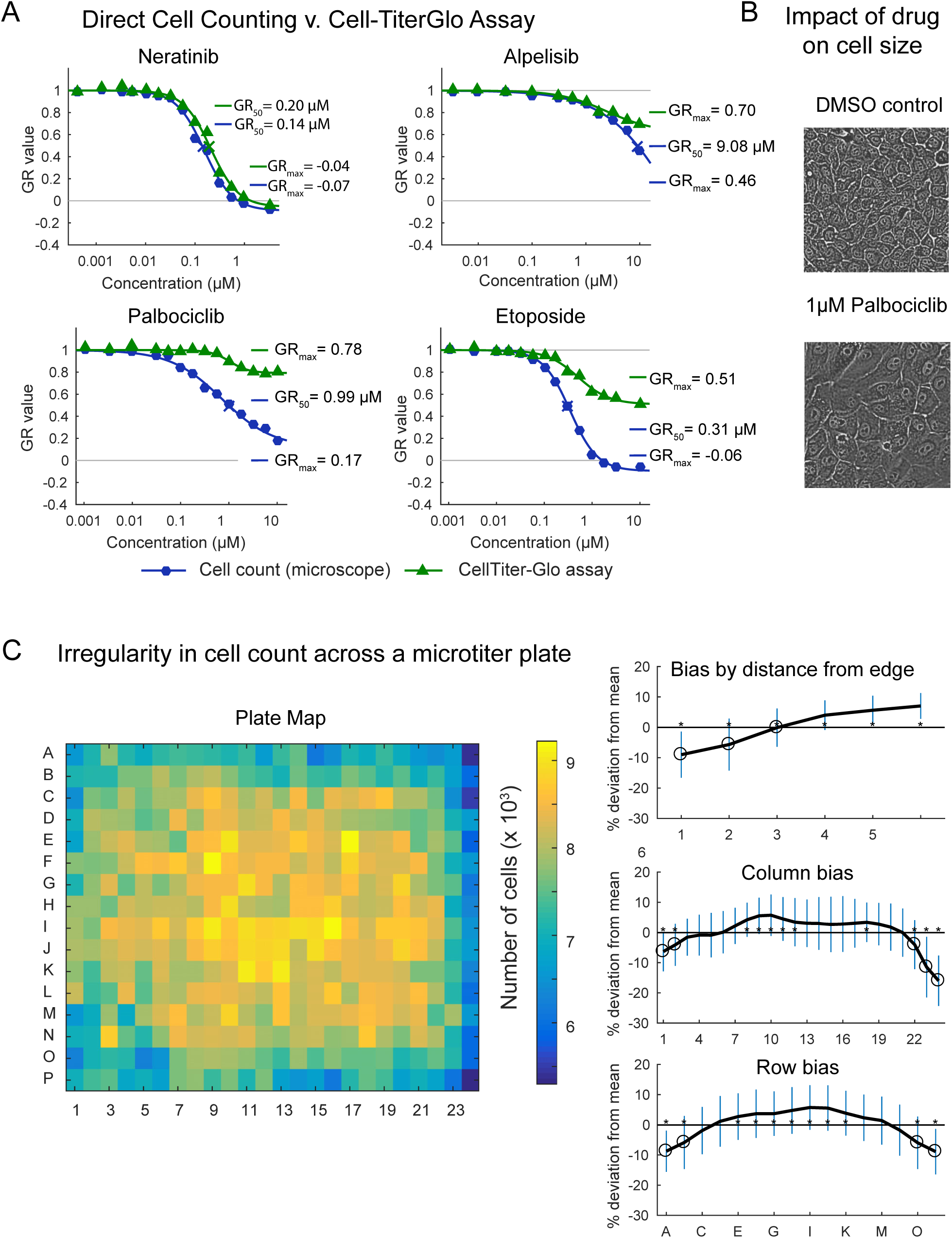
Experimental causes of variability. (A) Dose-response curves of MCF 10A cells treated with four different drugs measured by image-based cell count or ATP content (CellTiter-Glo®) on the same day by the HMS LINCS Center, which is equivalent to technical replicates. Note the GR_50_ value for alpelisib as measured by CellTiterGlo was not defined. (B) Representative images of MCF 10A cells treated with vehicle control (DMSO) or 1 µM Palbociclib. (C) Uneven growth of MCF 10A cells in a 384-well plate over the course of three days that demonstrates the presence of edge effects. In the heatmap, color represents the number of cells per well, as assessed by imaging. Plots show deviation from mean number (for the full plate based on the distance from the edge, by column, or by row. Error bars represent the standard deviation. Asterisks indicate the row or column differs significantly from all others.

It is known that CellTiter-Glo® and direct cell counts are poorly correlated when drugs cause dramatic changes in cell size or alter ATP metabolism, thereby changing the relationship between ATP level in a cell extract and viable cell number (Figure 2B for Palbociclib) (Harris et al., 2016a; Salani et al., 2013; Soliman et al., 2016). The magnitude of this effect depends on the drug being assayed and also on the cell line (Niepel et al., 2017); as a consequence, direct cell counting and CellTiter-Glo® can be substituted for each other in some cases but not in others. Thus, a substitution that appears to be justified by pilot studies on a limited number of cell lines and drugs can be problematic when the number and chemical diversity of drugs is increased. In this context, we note that counting viable cells by microscopy is both more direct and cheaper as a measure of viability than ATP levels; CellTiter-Glo® is used in place of counting primarily because it is perceived as being easier to perform. The problem is not with the CellTiter-Glo® itself, which is reproducible and can be precisely calibrated, but with equating reduced ATP levels with reduced cell counts. Situations in which ATP levels fall in viable or dividing cells might be of interest biologically but identifying these situations requires performing CellTiter-Glo® and cell counting assays in parallel.

Edge effects and non-uniform cell growth are a second substantial source of variation in cell based studies performed in microtiter plates (Bushway et al., 2010; Coyle et al., 1989) thought to arise from temperature gradients and uneven evaporation of media at the edges of plates. We have observed a variety of irregularities in plating and cell growth that often depend on the batch of microtiter plates, even when these plates are obtained from a single highly regarded vendor; we recommend testing all batches of plates for uniformity of cell growth (Niepel et al., 2017). A variety of approaches are available to minimize such effects (e.g. placing plates in humidified chambers to reduce evaporation from edge wells), but variation in growth is often confined to specific regions of a plate (Figure 2C) causing systematic errors in dose-response data. We have therefore found that randomized compound dispensing is valuable in mitigating biases introduced by edge effects and irregular growth. Using an automated liquid handling robot such as the HP D300e Digital Dispenser, it is possible to dispense compounds directly into microtiter plates in an arbitrary pattern, randomizing the locations of control and technical replicate samples in the plate. In this way, systematic error arising from edge effects is converted into random error, which is easily modeled statistically (Niepel et al., 2017). We have also found that the use of simple washing and dispensing robots reduces errors that humans make during repetitive pipetting operations. Most of these robots are small, robust, and relatively inexpensive, and our experience suggests that they greatly improve the reproducibility of medium-and high-throughput cell-based and biochemical studies.

A third source of error that we explored involves the concentration range over which a drug is assayed and the impact of the range on curve fitting and parameter estimation. For example, when we followed general practice and assayed Trametinib (a MEK kinase inhibitor) over a thousand-fold concentration range, growth of MCF 10A cells was fully arrested at ∼30 nM (Figure 3A, left plot): phenotypic response did not change even when the dose was increased 100-fold to 1 µM and thus, increasing the dose-range had no effect on curve fitting and parameter estimation (Figure 3A, left plot). However, when Dasatinib (a poly-selective SRC-family kinase inhibitor) was assayed over a thousand-fold range, curve fitting identified a plateau in GR value between 0.3 to 1 µM, but when the dose-range was extended to high drug concentrations GR values became negative, demonstrating a shift from cytostasis to cell killing (Figure 3A, right plot). Thus, a dose-range that is adequate for analysis of Trametinib is not adequate for Dasatinib. This sort of variation is difficult to identify in a high-throughput experiment and suggests that pilot empirical studies are needed to optimize dose ranges for specific compounds. Such variation did not impact reproducibility in our inter-Center study, because all Centers used the identical dose series, but dose range did affect the accuracy of GR_max_ estimation in general.

**Figure 3:**
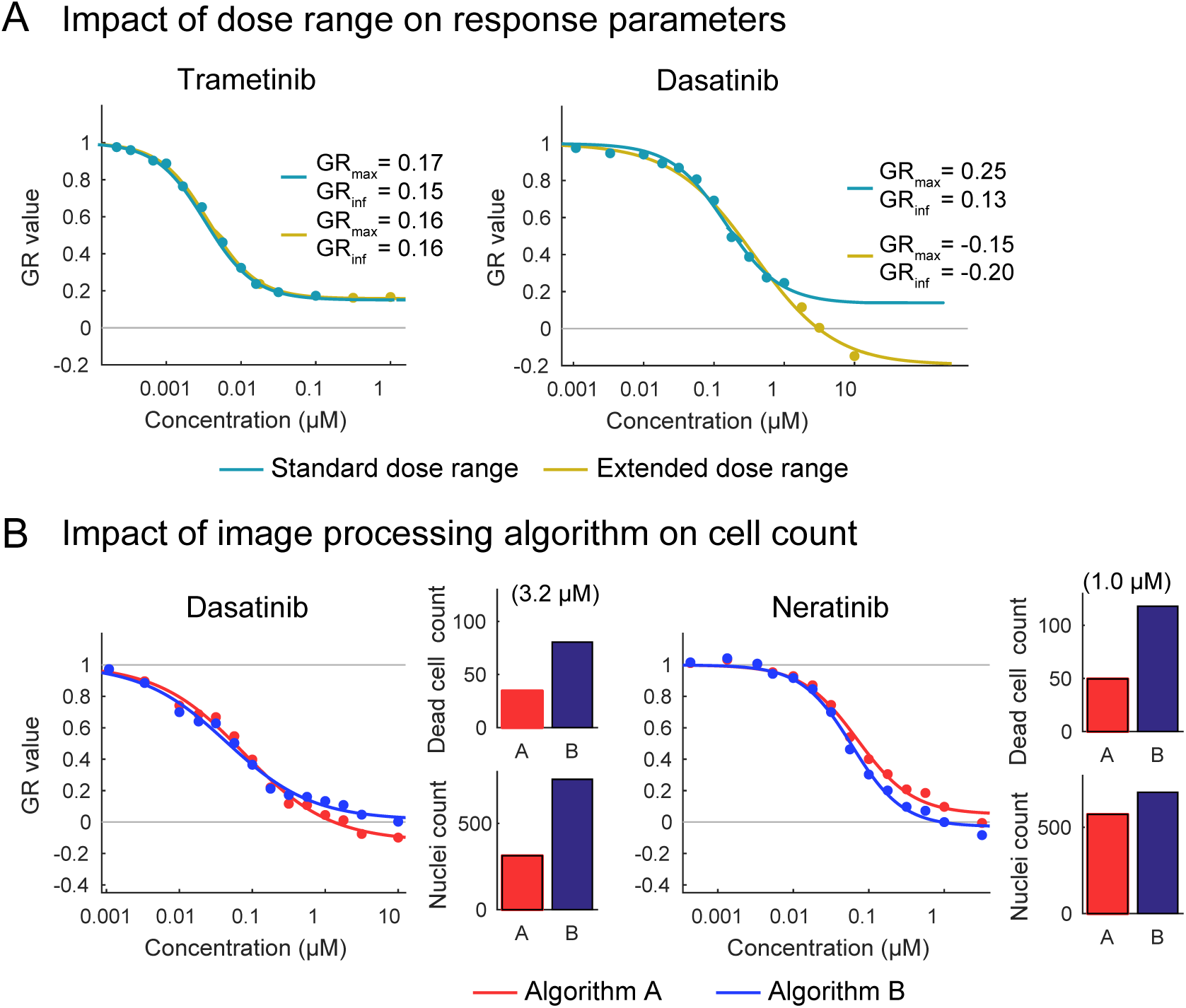
Technical causes of variability. (A) Dose-response curves of MCF 10A cells treated with Trametinib or Dasatinib fitted to either the extended dose range (up to 1 µM and 10 μM, respectively) or omitting the last order of magnitude. (B) Results of cell counting for MCF 10A cells treated with Dasatinib or Neratinib using two different image processing algorithms (denoted as A (red) and B (blue)) included in the Columbus image analysis software package. (C) Number of dead cells (LIVE/DEAD^™^ Fixable Red Dead Cell Stain positive) and nuclei (Hoechst positive) counted for MCF 10A cells treated with 3.16 µM Dasatinib or 1 µM Neratinib based on the two different algorithms (corresponding to the plots in C).

A fourth source of inter-Center variation was apparent for Centers that used imaging-based cell counting. It involved differences in cell counts for MCF 10A cells treated with high doses of Dasatinib and Neratinib (Figure 3B). Above 1 µM, GR values were reproducibly negative at LINCS Center One for both drugs but in one LINCS Center GR_max_ was consistently above 0. Follow-up studies showed that the discrepancy arose from the use of image processing algorithms that included dead cells in the “viable cell count” and from over-counting the number of cells when multi-nucleation occurred (Orth et al., 2011; Röyttä et al., 1987). Observed differences in drug response across centers could be recapitulated in a single laboratory by using different image processing routines and were also evident by visual inspection of the segmented images (Figure 3B). In retrospect, all Centers should have processed images in the same way using Dockerized software (List, 2017), but the necessary routines are often built into manufacturer’s proprietary software, making standardization of image analysis dependent on the availability of primary data. This demonstrates the importance of locking down all steps in the data processing pipeline from raw measurements to final parameter estimation, as well as a relatively subtle interplay between biological and technical sources of variability. Unfortunately, such harmonization is difficult to achieve when replicating published results that do not provide primary data.

### Biological factors impacting repeatability

Variables that can change the biology of drug response, such as media composition, incubation conditions, microenvironment, media volume, and cell density, have been discussed elsewhere (Hafner et al., 2016; Haverty et al., 2016) and were controlled to the greatest extent possible in this study through standardization of reagents and the use of GR metrics. In a truly independent repeat of the current study, experimental variables such as these would need to be considered as additional confounders, because it is difficult to fully standardize a reagent as complex as tissue culture media. However, one center performed a preliminary comparison of batches of horse serum, hydrocortisone, cholera toxin, and insulin and found that the effects on drug response were smaller than the sources of variation discussed above (data not shown).

At the outset of the study, we had anticipated that the origin of the MCF 10A isolate would be an important determinant of drug response. MCF 10A cells have been grown for many years, and karyotyping reveals differences among isolates (Caruso et al., 2001; Cowell et al., 2005; Kim et al., 2008; Marella et al., 2009; Soule et al., 1990; Zientek-Targosz et al., 2008), which is why we had distributed aliquots of a single isolate to all Centers). To investigate the potential impact of genetic drift, we assembled MCF 10A isolates from different laboratories and Center One then compared them to each other and to a histone H2B-mCherry-tagged subclone of one of the isolates (Figure S4A); we also examined four subclones from the LINCS MCF 10A master stock. Variation in measured drug response across all of these isolates and subclones was substantially smaller than what was observed when a single isolate was assayed at different Centers. GR metrics correct for differences in growth rates but we nonetheless compared doubling time across Centers; we found them to be highly consistent among those Centers using cell counting assays and slightly higher when CellTiterGlo was used to estimate cell number (Figure S4B). We conclude that even though clonal variation can have a substantial variation on drug response and other properties of cultured cells (Ramirez et al., 2016), it was not a significant contributor to variation in dose-response data in this study.

The duration of drug exposure is not generally explored in *in vitro* studies, and instead assays are usually performed a fixed time after treatment. To investigate the impact of time of drug exposure we monitored the responses of MCF 10A cells to drugs in a live-cell experiment in which cell number was measured every two hours using an automated high-throughput microscope. We then quantified the response by calculating GR values over a 12-hour moving window (yielding time-dependent GR values) and found that the effect of time and dose were substantial in some cases but not others. For example, GR values for cells exposed to Etoposide were nearly constant across all doses throughout a 50-hour assay period (Figure 4, top left plot), whereas GR values for Neratinib varied from 0 to 1 over the same period (Figure 4, bottom left plot), with the highest variability at intermediate drug doses. As a consequence, GR dose-response curves and metrics derived from these curves, such as GR_50_ and GR_max_, varied with time (Figures 4 and S5). This observation again emphasizes the interplay between biological factors and metrics of drug response. The temporal dependence of drug response is likely to reflect biological adaptation, drug export, and other factors important in drug mechanism of action (Fallahi-Sichani et al., 2017; Fletcher et al., 2010; Hafner et al., 2016; Harris et al., 2016b; Muranen et al., 2012). These factors remain largely unexplored and are likely to add substantially to the difficulty of reproducing experimental data when protocols are not carefully followed.

**Figure 4:**
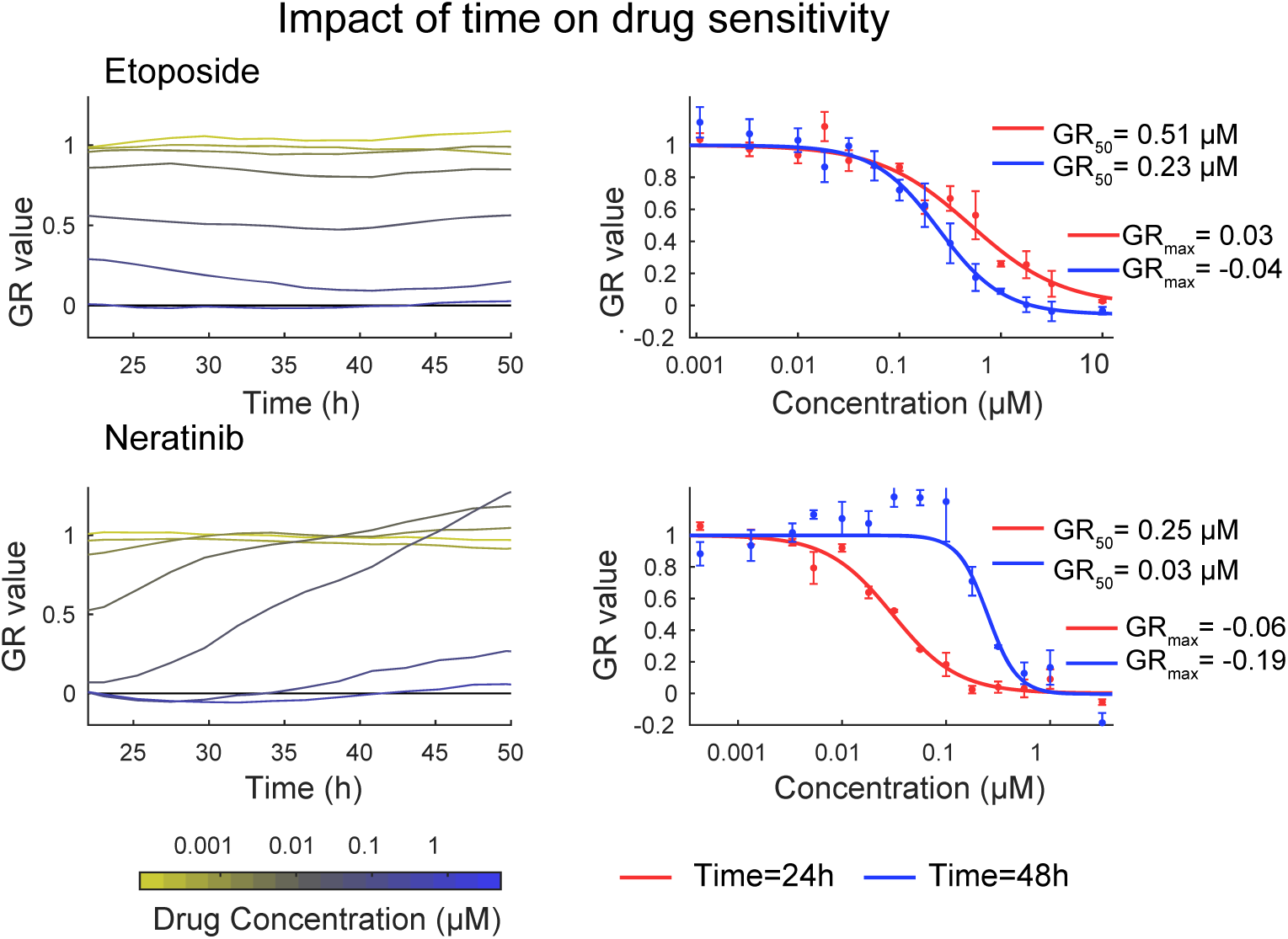
Changes in drug response related to the underlying biology. Left panels: Inhibition of MCF 10A growth (12-hour instantaneous GR values) measured in a time-lapse, live-cell experiment involving treatment with multiple doses of Etoposide (top) or Neratinib (bottom). Different colors indicate different drug concentrations ranging from 1 nM (yellow) to 10 µM (blue). Right panels: Dose-response curves derived from 12-hour GR values computed at 24 (red) and 48 hours (blue) across three biological repeats. Etoposide displays only modest time-dependent effects while neratinib appears to be more effective at inhibiting growth at early time points as compared to later time points.

**Figure 5:**
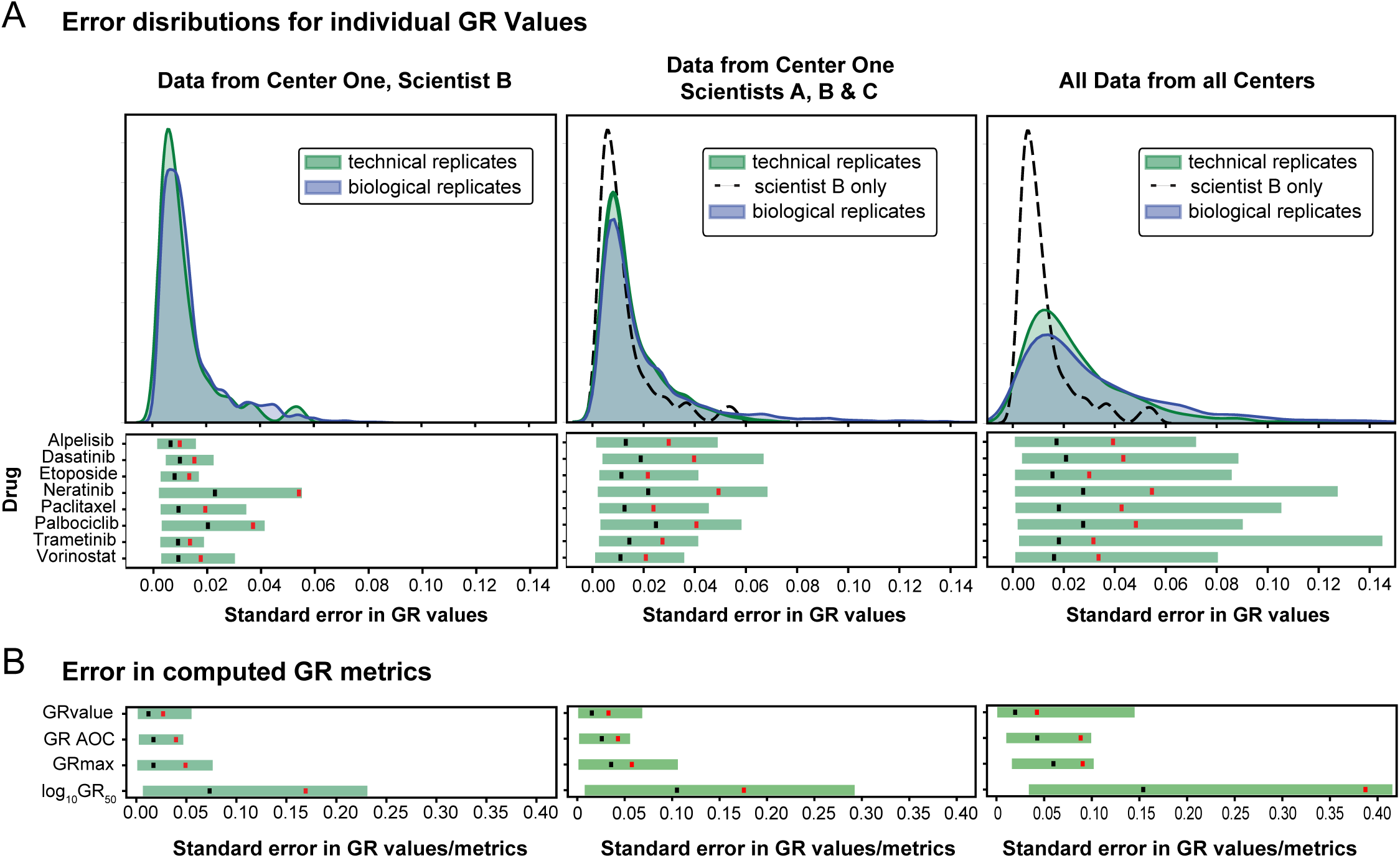
Technical and biological variability in estimating GR values and metrics. (A) The distribution of standard error in the measurement of GR values across technical (green curve) or biological (blue curve) replicates for all drug and dose points. The left panel depicts data from Center One, scientist B (performed in 2018); the middle panel four sets of measurements from all scientists in Center One (performed between 2016-2018); and the right panel all data from all Centers. The distribution of technical error for Scientist B is duplicated in the middle and right panels as a black dotted line. Data for these distributions derive from GR values, not GR metrics, and in the case of the left panel for example, involve 576 data points (8 doses × 8 drugs × 3 biological repeats × 3 technical repeats). The lower section of each panel depicts the error in GR value measurements across technical replicates for each individual drug. (B) The range of standard error in GR values compared to the standard error in corresponding GR metrics (GR_max_, area over the GR curve (GR AOC), and log10GR50) for all drugs. The black vertical line (A, lower plots, and B) is the mean error for a given drug and the red vertical line demarcates the 90^th^ percentile error (i.e the error for 90% of GR values or metrics is below that value).

### Final results

To determine how successful we had been in identifying and controlling for sources of variability in the measurement of drug dose-response, we performed two sets of tests. First, measurements for all drugs were repeated in LINCS Center One two years after the first round of studies by an experienced research scientist (Scientists A from the original study, Figures 1 and S2) and by a newly recruited technical associate (Scientist B) who did not have prior experience with GR metrics. Data were collected in biological triplicate with each replicate separated by a minimum of one cell passage from the next; each biological replicate was assayed in technical triplicate, as described in Figure 1B. Plates, media, supplements and serum were all from different batches as experiments performed in 2017 and cells were recovered from independent frozen stocks. However, the protocol remained the same over the two-year period and involved the same automated compound dispensing and plate washing procedures.

Data from newly trained Scientist B exhibited similar standard error for biological and technical repeats with a standard error for estimation of GR values of 0.012 across all drugs, doses and repeats. The distribution was long tailed, an apparent consequence of systematic error in assays involving Neratinib (Figure 5A; lower panel). As shown in Figure 4, GR values for Neratinib are strongly time-dependent and we might therefore expect data for this drug to be sensitive to small variations in procedure. The observed error in GR values corresponds to a difference in the estimation of GR50 values of 1.17-fold (mean standard error, which corresponds to a variation of ± 0.07 in log_10_(GR_50_)) while the standard error for 90% of GR50 values corresponded to a difference ∼1.5-fold (± 0.18 in log_10_(GR_50_)) (Figure 5B). For all measurements obtained in Center One over a period of two years, the mean standard error in GR values was 0.015 only slightly higher than the error from Scientist B alone. The standard deviations in log_10_(GR_50_) and GR_max_ values obtained by Scientist A over a two year period were indistinguishable and there was no observable batch effect for any drug (Figure S2). These distributions represent our best estimate of the error associated with measuring drug-dose response using a single protocol and experimental setup but different consumables; this error estimate can therefore be incorporated into future error models. In our opinion these values also represent a good level of accuracy and reproducibility.

As a second test, all LINCS Centers repeated drug-dose response measurements using their closest approximation to the standard protocol. One Center used CellTiter-Glo® rather than direct cell counting to estimate viable cell number. Use of this method resulted in greater deviation from the results in Center One, as expected from the studies shown in Figures 2-6 (e.g. technical error in the CellTiter-Glo® data from Center Four exceeded that of all other centers). Despite such differences in procedure inter-Center variability at the end of the study was dramatically lower than at the outset, with a standard error in GR value measurement ∼2-fold higher than in Center One and errors in the estimation of GR_50_ of ∼2 standard deviations. The mean standard error for log_10_(GR_50_) across all drugs was ± 0.15 while the standard error for 90% of measured GR50 values was within ∼2.5-fold (±0.38 in log_10_(GR_50_)) (Figure 5B). The largest identifiable source of error in the final data arose from use of the CTG assay (Figure 6).

**Figure 6:**
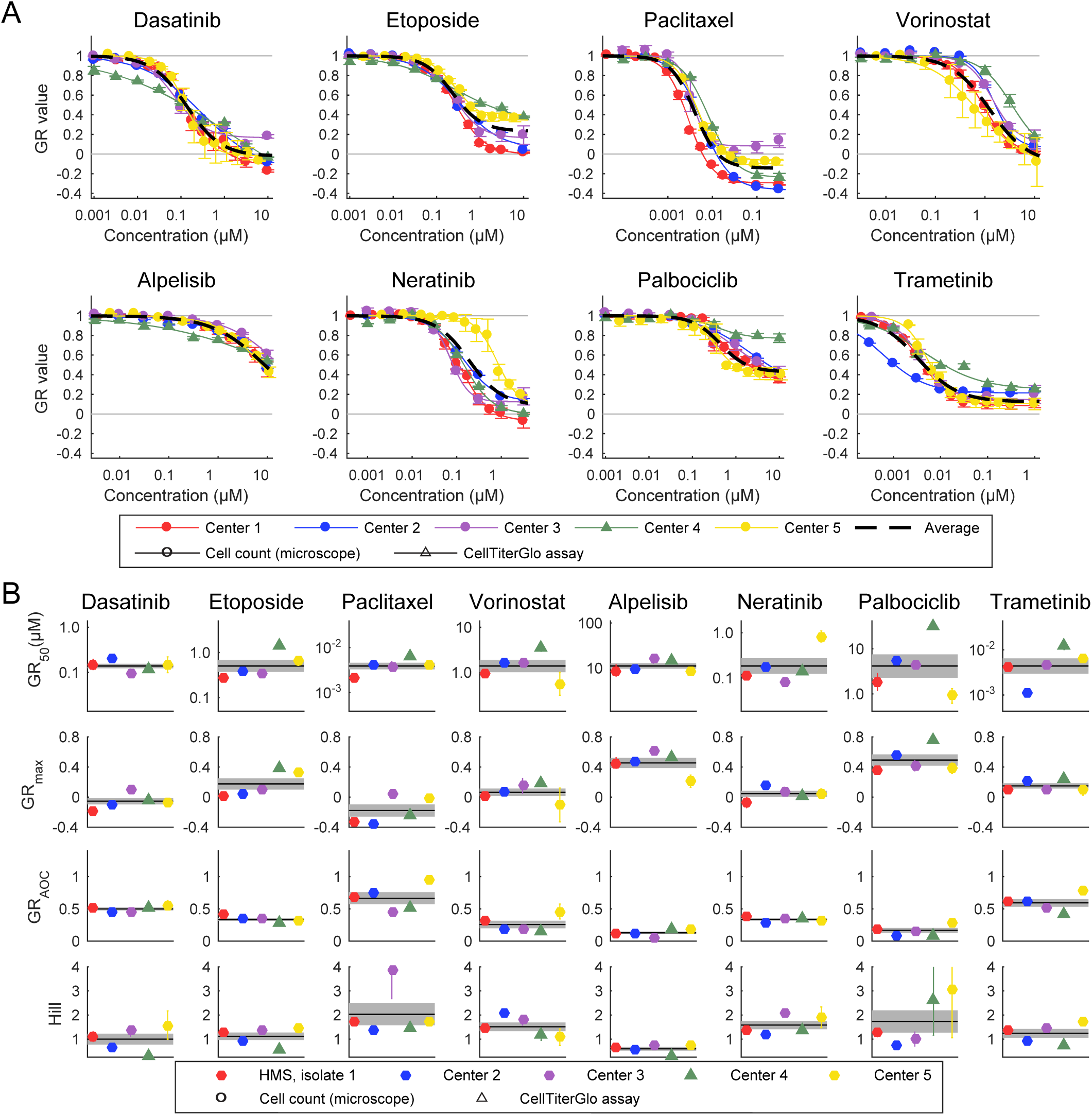
Variability of the response measures across Centers. (A) Dose-response curves of MCF 10A cells treated with eight drugs measured independently by the five LINCS Centers (circles represent data from image-based assays and triangles from CellTiter-Glo® assays). See Figure S6 for underlying replicates. Dotted black lines show the dose-response curve when all independent replicates were averaged. (B) GR metrics describing the sensitivity of MCF 10A cells to eight drugs measured independently by five LINCS Centers (circles represent data from image-based assays and triangles from CellTiter-Glo® assays). The black line shows the mean sensitivity across all Centers, and the gray area shows the standard error of the mean computed from the average of each center. For GR_50_ and GR_max_, error bars represent the standard deviation of the log10(GR) values. (C) Differences in repeatability/precision vs. biological sensitivity/stability/robustness. Note that some data are shared between Figures 6 and S3.

From these data we conclude that it is possible for previously inexperienced individuals to measure drug-dose response with high reliability over an extended period of time and that multiple Centers can approximate this level of reproducibility. However, deviations from an SOP with respect to automation and type of assay, which might be necessary for practical reasons, have a substantial negative impact.

## DISCUSSION

The observation that a significant fraction of biomedical research cannot be reproduced is troubling; it handicaps academic and industrial researchers alike and has generated extensive comment in the scientific and popular press (Arrowsmith, 2011; Baker, 2016; Begley and Ellis, 2012; Prinz et al., 2011; Wilkinson et al., 2016). The key question is why such irreproducibility arises and how it can be overcome; in the absence of such studies, FAIR data will remain little more than an aspiration. In this study we investigated the precision and reproducibility of a prototypical perturbational experiment performed in cell lines: drug dose-response as measured by cell viability. Perturbational experiments are foundational in genetics, chemical biology, and biochemistry and, when they involve human therapeutics, they are also of translational value. A consortium of five geographically dispersed NIH LINCS Centers initially encountered high levels of inter-Center variability in estimating drug-potency, even when a common set of reagents was used. Subsequent study in a single center uncovered possible sources of measurement error, resulting in a substantial increase in inter-Center reproducibility. Nonetheless, the final level of inter-center variability exceeded what could be achieved in a single laboratory over a period of two years by three different scientists. We ascribe the remaining irreproducibility to differences in compound handling, pipetting, and cell counting that were not harmonized because of the expense of acquiring the necessary instrumentation and a belief—belied by the final analysis—that counting cells is such a simple procedure that different assays can be substituted for each other without consequence. We believe the final level of intra-and inter-Center precision to exceed the norm for this class of experiments in the current literature (although this is not easy to prove) and that it provides a roadmap for future experiments of this type.

At the outset of the study we had hoped that comparison of data across Centers would serve to identify the specific biological, experimental, and computational factors that had the largest impact on data reproducibility. However, we discovered that most examples of irreproducibility were themselves irreproducible and that technical factors responsible for any specific outlier measurement were difficult to pin down. We therefore undertook a systematic study of the assay itself, in a single Center, with an eye to identifying those variables with the greatest impact on reproducibility. We found that these variables differed from what we expected *a priori*. For example, isolate-to-isolate differences in MCF 10A cultures had substantially less of an effect on drug response assays (Figure S4A) than the ways in which drugs and cells were plated into multi-well plates and counted (Figures 2-3).

In general, we found that irreproducibility most commonly arose from unexpected interplay between experimental protocol and true biological variability. For example, estimating cell number from ATP levels using the CellTiter-Glo® assay produces very similar results to direct cell counting with a microscope in the case of Neratinib, but this is not true for Etoposide or Palbociclib (Figure 2A). The discrepancy most likely arises because ATP levels in lysates of drug-treated cells vary for reasons other than loss of viability; these include changes in cell size and metabolism. We have previously shown that the density at which cells are assayed also has a dramatic effect on drug response (Hafner et al., 2016), but this too is context dependent. For some cell line-drug pairs, density has little or no effect, whereas for other pairs it increases drug sensitivity and for yet others it has the opposite effect. This observation has important implications for the design of experiments in which diverse compounds are screened: pilot studies on a limited range of conditions (dose and drug identity in this work) cannot necessarily be extrapolated to large datasets and are not a sound basis for substituting indirect assays for direct assays. The tendency for even experienced investigators to substitute assays for each, or to stick to historical methods rather than standardized protocols (SOPs), is undoubtedly a source of irreproducibility.

Several lines of evidence suggest that context dependence in drug response reflects real changes in the underlying biology and not flaws in assay methodology itself. For example, cell density directly impacts media conditioning and the strength of autocrine signaling, which in turn changes responsiveness to some drugs but not others (Wilson et al., 2012; Yonesaka et al., 2008). Thus, even in cell lines, drug-response is not a simple biological process and changes in measurement procedure that might have no effect in one cell type or biological setting can substantially affect results obtained in other settings. At the current state of knowledge, there is no substitute for empirical studies that carefully assess the range of conditions over which data remain reliable and precise for cell lines and drugs of interest. Moreover, the most direct assay - not a convenient substitute - should be used to score a phenotype whenever possible. Unfortunately, when the goal is collection of a large dataset, a prerequisite for most machine learning approaches, attention to biological factors known to be important from conventional cell biology studies is often deemphasized in favor of throughput.

Data processing routines are important for reproducibility (Sandve et al., 2013). Data and data analysis routines can interact in multiple ways, some of which are clear in retrospect but not necessarily anticipated. For example, collecting 8-point dose response curves generally represents good practice, but it is essential that the dose range effectively span the GEC_50_ (the mid-point of the response). When this is not the case (as illustrated by Figure 3A), curve fitting is underdetermined and response metrics become unreliable. In many cases problems with dose range are not evident until an initial assay has been performed and an iterative approach is necessary. Iteration is straightforward in small-scale studies, but substantially harder for large-scale screens; for a large dataset, data processing routines should automatically identify and flag problems with dose range. Additionally, accurate reporting of dose range is necessary to provide a bound to drug sensitivity measurement. Another example involves image processing routines for automated cell counting: such routines should be optimized for cells that grow and respond to drugs in different ways (Figure 3B) and must be tested for performance at high and low cell densities.

Processing pipelines for the type of data collected in this study are much less developed than the pipelines commonly used for genomics data (Ashley, 2016; Bao et al., 2014; Lam et al., 2011), but much can be learned from the comparison. For example, computational platforms with provenance such as Galaxy (Goecks et al., 2010), Sage Bionetworks’ Synapse (Omberg et al., 2013), or Cancer Genomics Clouds, have been developed to support data sharing, reproducible analyses, and transparent pipelines, with a primary focus on genomics data. Galaxy also provides a shared platform on which to execute workflows, which serves to eliminate compute environment differences. With sufficient effort dedicated to pipeline development it should be possible to adapt such solutions to a wider range of assay types. In the specific case of dose-response data we have developed an on-line set of Jupyter notebooks (https://github.com/labsyspharm/MCF10A_DR_reproducibility and https://github.com/datarail/datrail) and a list of best practices at http://www.grcalculator.org (Clark et al., 2017). Image processing algorithms present a unique challenge in that they are frequently embedded in proprietary software linked to a specific data acquisition microscope, which complicates common analysis across laboratories; publicly available (and often open-source) image analysis platforms are always preferable (Carpenter et al., 2006).

### Elements of a reproducible workflow

The elements of a workflow for reproducible collection of dose-response data are fairly simple and outlined in Figure 7A: (i) *Standardization of reagents* including obtaining cell lines directly from repositories such as the ATCC, performing mass spectrometry-based quality control of small molecule drugs, and tracking lot numbers for all media additives; (ii) *Standardized data processing* starting with raw data and metadata through to reporting of final results; and (iii) *Use of automation* to improve reliability and enable experimental designs too complex or labor intensive for humans to execute reliably; in many cases this involve simple and relatively inexpensive bench-top dispensing and washing (iv) *Close attention to metrology* (analytical chemistry), measurement technology and internal quality controls. The first two points are obvious, but not trivial to implement, because laboratories are not all equipped the same way and some data processing routines are embedded in a non-obvious way in instrument software. In the current work, a major benefit of automation is that it makes random plate layouts feasible, thereby changing systematic edge effects into random error that has a reduced impact on the dose-response measures. In the case of dose-response data, metrology focuses on variability among technical and biological replicates, assessment of edge effects, and outlier detection. Edge effects and other spatial artifacts can be identified by statistical analysis (Mazoure et al., 2017) and plate-wise data visualization (Boutros et al., 2006). Spatial artifact can then be removed with plate-level normalization such as LOESS/LOWESS smoothing (Boutros et al., 2006; Pelz et al., 2010), spatial autocorrelation (Lachmann et al., 2016), or statistical modeling (Mazoure et al., 2017).

**Figure 7:**
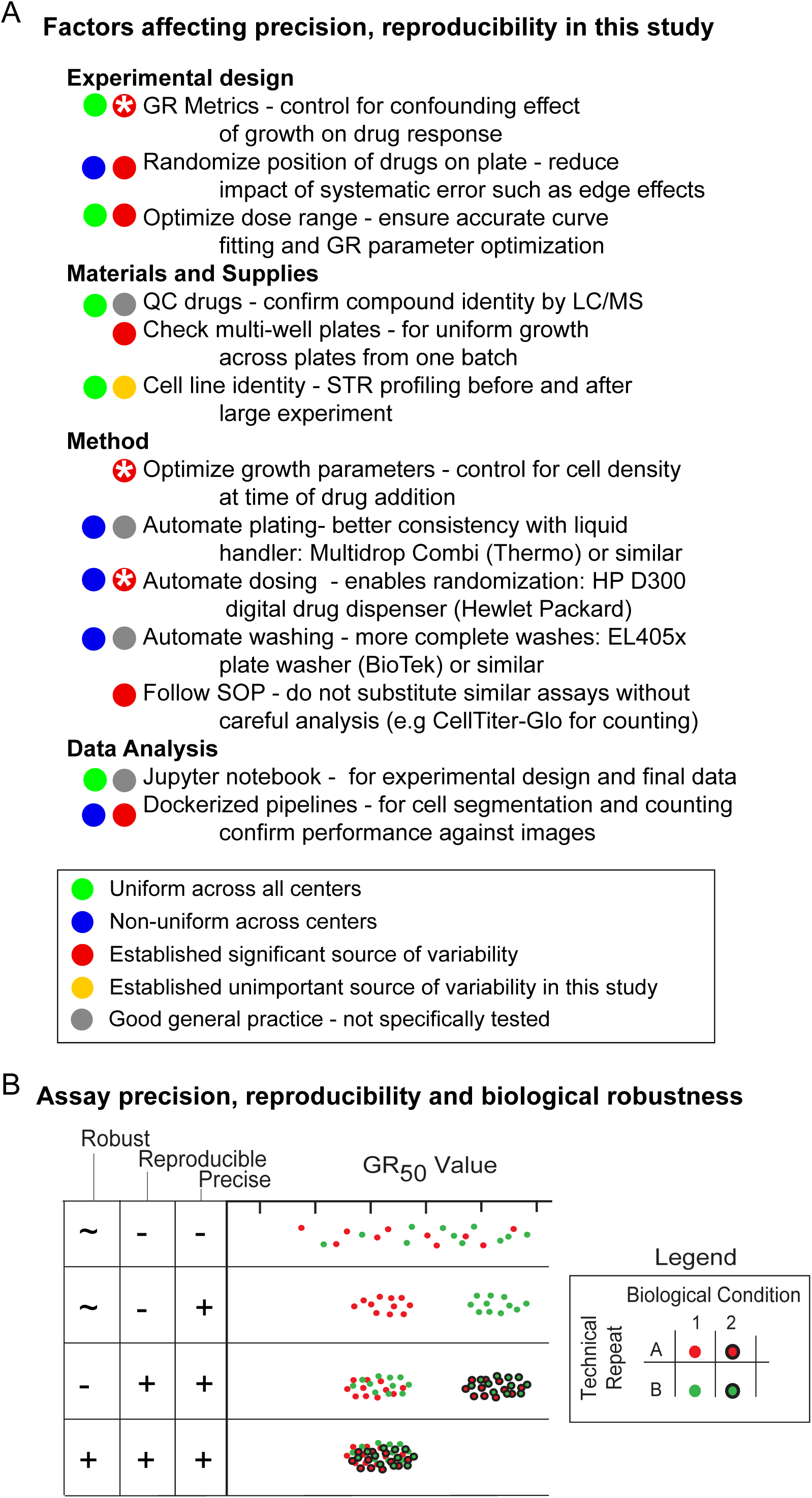
Best practices for dose response measurement experiments. (A) Summary of findings in this and related studied with respect to experimental and technical variability in dose response studies at the experimental design, materials, methods, and analysis stages; “*” indicates sources of variability that have been thoroughly investigated in a previous paper (Hafner et al., 2016). Jupyter notebooks for experimental design and data analysis are available: https://github.com/labsyspharm/MCF10A_DR_reproducibility and https://github.com/datarail/datrail. (B) Differences between precision, robustness and reproducibility; see text for details.

The primary contribution of the current study is to show that future execution of reproducible drug dose-response assays in different cell types requires systematic experimentation aimed at establishing the robustness of assays over a full range of biological settings and cell types. Such robustness is distinct from conventional measures of assay performance such as precision or repeatability in a single biological setting (Figure 7B). Testing of this type is not routinely performed for the simple reason that establishing and maintaining robust and reproducible assays is time consuming and expensive: we estimate that reproducibility adds ∼20% to the total cost of a large-scale study such as drug-response experiments in panels of cell lines (AlQuraishi and Sorger, 2016). Iterative experimental design is also essential, even though it has been argued that this is not feasible for large-scale studies (Harris, 2017).

## Conclusions

A question raised by our analysis is whether, given their variability and context-dependence, drug response assays performed *in vitro* are useful for understanding drug response in other settings, human patients in particular (Wilding and Bodmer, 2014). Concern about the translatability of *in vitro* experiments is long-standing, but we think the current work provides grounds for optimism rather than additional worry. Simply put, if *in vitro* data cannot be reproduced from one laboratory to the next, then it is no wonder that they cannot easily be reproduced in humans; conversely, paying greater attention to accurate and reproducible *in vitro* data is likely to improve translation. Many of the factors that appear to represent irreproducibility in fact arise from biologically meaningful variation. This includes the time-dependence of drug response, the impact of non-genetic heterogeneity at a single-cell level, and the influence of growth conditions and environmental factors (Cohen et al., 2008; Loewer and Lahav, 2011; Muranen et al., 2012; Wilson et al., 2012; Yonesaka et al., 2008). The simple assays of drug response in current use are unable to correct for such variability, and the problem is made worse by “kit-based science” in which technical validation of assays is left to vendors. However, if the challenge of understanding biological variability at a mechanistic level is embraced, it seems likely that we will improve our ability to conduct *in vitro* assays reproducibly and apply data obtained in cell lines to human patients. We note that RNAi, CRISPR and other perturbational experiments in which phenotypes are measured in cell culture are likely to involve many of the same variables as the dose-response experiments studied here.

Despite a push for adherence to the FAIR principles (Wilkinson et al., 2016) there is currently no consensus that the necessary investment is worthwhile, nor do incentives exist in the publication or funding processes for individual research scientists to meet FAIR standards (AlQuraishi and Sorger, 2016; Goodspeed et al., 2016). Data repositories are essential, but we also require better training in metrology, analytical chemistry and statistical quality control. In developing incentives and training programs, we must also recognize that reproducible research is a public good whose costs are borne by individual investigators and whose benefits are conferred to the community as a whole.

## MATERIAL AND METHODS

### Cell lines and drugs

Three isolates of MCF 10A, here referred to as MCF 10A-GM, MCF 10A-OHSU, and MCF 10A-HMS, were sourced independently at three different times from the ATCC. MCF 10A-H2B- mCherry cells were created by inserting an H2B-mCherry expression cassette into the AAVS1 safe harbor genomic locus of MCF 10A-HMS using CRISPR/Cas9 (Hafner et al., 2016). All lines were confirmed to be MCF 10A cells by STR profiling (Table S1), and confirmed to have stable karyotypes by g-banding 47,XX,i(1)(q10),+del(1)(q12q32),add(3)(p13),add(8)(p23),add(9)p(14). All lines were cultured in DMEM/F12 base media (Invitrogen #11330-032) supplemented with 5% horse serum (Sigma-Aldrich #H1138), 0.5 μg/mL hydrocortisone (Sigma # H-4001), 20 ng/mL rhEGF (R&D Systems #236-EG), 10 μg/mL insulin (Sigma #I9278), 100 ng/mL cholera toxin (Sigma-Aldrich #C8052), and 100 units/mL penicillin and 100 μg/mL streptomycin (Invitrogen #15140148 or #15140122 or other sources) as described previously (Debnath et al., 2003). Base media, horse serum, hydrocortisone, rhEGF, insulin, and cholera toxin where purchased by the MEP-LINCS Center and distributed to the remaining experimental sites. MCF 10A-GM was expanded by Gordon Mills at MD Anderson Cancer Center and distributed to all experimental sites. Cell identity was confirmed at individual experimental sites by short tandem repeat (STR) profiling, and the cells were found to be free of mycoplasma prior to performing experiments.

Drugs were obtained from commercial vendors by HMS LINCS, tested for identity and purity in house as described in detail in the drug collection section of the HMS LINCS Database (http://lincs.hms.harvard.edu/db/sm/), and distributed as 10 mM stock solutions dissolved in DMSO to all experimental sites. See Table S2 for additional metadata for key reagents.

### Drug response experiments and data analysis

The experimental and computational protocols to measure drug response are described in detail in two prior publications (Hafner et al., 2017b; Niepel et al., 2017). The following protocol was suggested for this study: cells were plated at 750 cells per well in 60 µL of media in 384-well plates using automated plate fillers and incubated for 24 h prior to drug addition. Drugs were added at the indicated doses with a D300 Digital Dispenser (Hewlett-Packard), and cells were further incubated for 72 h. At the time of drug addition and at the endpoint of the experiment cells were staining with Hoechst and LIVE/DEAD™ Fixable Red Dead Cell Stain (ThermoFisher Scientific) and cell numbers were determined by imaging as described (Hafner et al., 2016; Niepel et al., 2017) or by the CellTiter-Glo® assay (Promega). Some details of the experimental protocol differed across Centers and over time, e.g. manually dispensing of drugs or use of 96-well plates. The data included in Figures 1C, and S2 (Scientist C) were collected for a separate project, and included here as an additional comparison. In these experiments, cells were treated via pin transfer, and HMS isolate 3 MCF10A cells were used.

For live-cell experiments with MCF 10A-H2B-mCherry, cell counts were performed by imaging plates in an 2 hr interval over the course of 96 hours (only first 50 hours shown) (Hafner et al., 2016; Niepel et al., 2017). Data analysis was performed as described previously (Hafner et al., 2016; Niepel et al., 2017).

Irregularities in growth across microtiter plates was performed by plating MCF 10A cells at 750 cells per well in 60 µL of media in 384-well plates using automated plate fillers and determining cell numbers after 96 h through imaging as described (Hafner et al., 2016; Niepel et al., 2017).

## DECLARATIONS

### Acknowledgements

We thank K. Ward for arranging the HP D300 instrument trial at Mount Sinai.

### Authors’ contributions

M.N., L.M.H., M.R.B., C.E.M. and E.H.W. designed the study and led its execution; P.K.S supervised the systematic analysis of assay variability and oversaw manuscript preparation. M.H. contributed to experimental design and M.H and K.S performed all data analysis. A.M.B. and A.D.S. participated in data collection and initial processing. All authors participated in editing the manuscript.

### Materials & Correspondence

Data presented in this paper are included in the additional material and available on the GR browser at http://www.grcalculator.org/grbrowser/.

---

***Figure S1: GR dose-response curve and metrics***

Schematic of a dose-response curve under the GR model and the source of the derived metrics.

***Figure S2: Assay stability over time***

GR metrics of MCF 10A cells treated with each drug showing the mean and standard deviation of biological triplicates collected by an experienced research scientist (Scientist A, circles), by the same scientist two years later (squares), by a new technician two years later (Scientist B, triangles), and biological duplicates collected as part of a separate LINCS project in Center One (http://lincs.hms.harvard.edu/db/datasets/20344/) (Scientist C).

***Figure S3: GR metrics describing the initial experiments to assess sensitivity of MCF 10A cells to eight drugs measured at three Centers.***

Center Three and Center Four represent the final results provided by each Center and are the same data presented in Figure 6A. Preliminary 1 represents initial experiments run by Center Three, which showed poor agreement with data from the other two Centers (see table to pipeline details). The disparate results reflect in part differences in readout (CTG vs. Imaging) for Etoposide, Vorinostat, Alpelisib, and Palbociclib. Preliminary 2 represents a coordinated effort by Centers Three and Four: daughter drug plates used by Center Three in Preliminary 1 experiments were shipped to Center Four for cell culture. For many drugs, there is good agreement between Preliminary 1 and Preliminary 2, indicating consistency between cell culturing. The consistent discrepancy in responses to Trametinib between Preliminary 1-Preliminary 2 studies and Center Three-Center Four studies indicate errors in construction of the drug dilution series.

***Figure S4: GR metrics across MCF 10A isolates***

(A) GR metrics describing the sensitivity of four different MCF 10A cell isolates to eight drugs measured independently at the HMS LINCS center. The black line shows the mean sensitivity measured across all isolates, and the gray box shows the standard error of the mean. (B) The doubling time of MCF10A cells at each LINCS Center. The error bars represent the standard deviation of the mean, the * indicates P < 0.05 by one way ANOVA with Tukey’s multiple comparison tests.

***Figure S5: Instantaneous GR metrics***

Instantaneous GR_max_ (A) and GR_50_ (B) values in MCF 10A cells treated with Etoposide (left) and Neratinib (right) over the course of 24 hr for three biological repeats.

***Figure S6: Technical and biological variability in GR metrics across Centers***

GR metrics for all technical and biological replicates of MCF 10A cells treated with eight drugs by five LINCS Centers (circles represent data from image-based assays and triangles from CellTiter-Glo® assays).

## ADDITIONAL MATERIAL

### Supplemental Data 1

Final drug-response data generated by all LINCS Centers (Figure 5).

### Supplemental Data 2

Data generated by the HMS LINCS Center during follow-up experiments (Figures 1-4 and S2, S4-S5).

### Supplemental Data 3

All technical and biological replicate data generated by all Centers (Figure S6).

### Table S1: STR profiles

Results of STR profiling on all MCF 10A isolates, and clones used in this work.

### Table S2: Key reagent metadata

Source, catalogue, and lot numbers for all media components and small molecules used in this work.

