## Supplemental Information for "A multi-center study on factors influencing the reproducibility of *in vitro* drug-response studies"

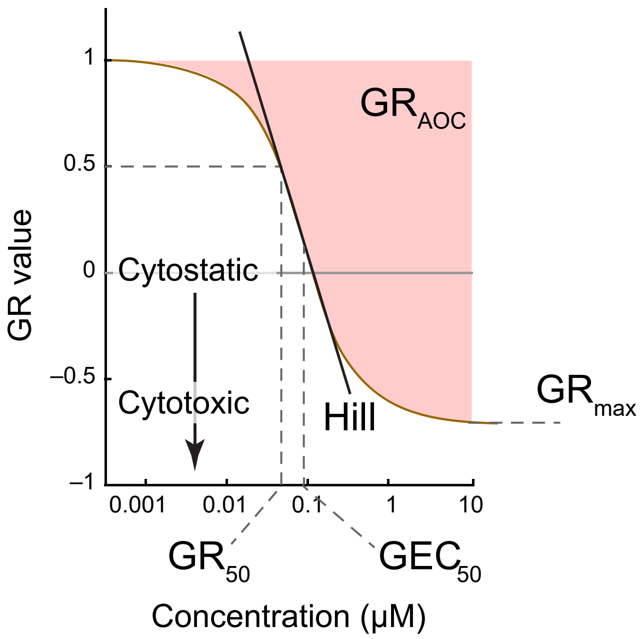

Figure S1, Related to Figure 1

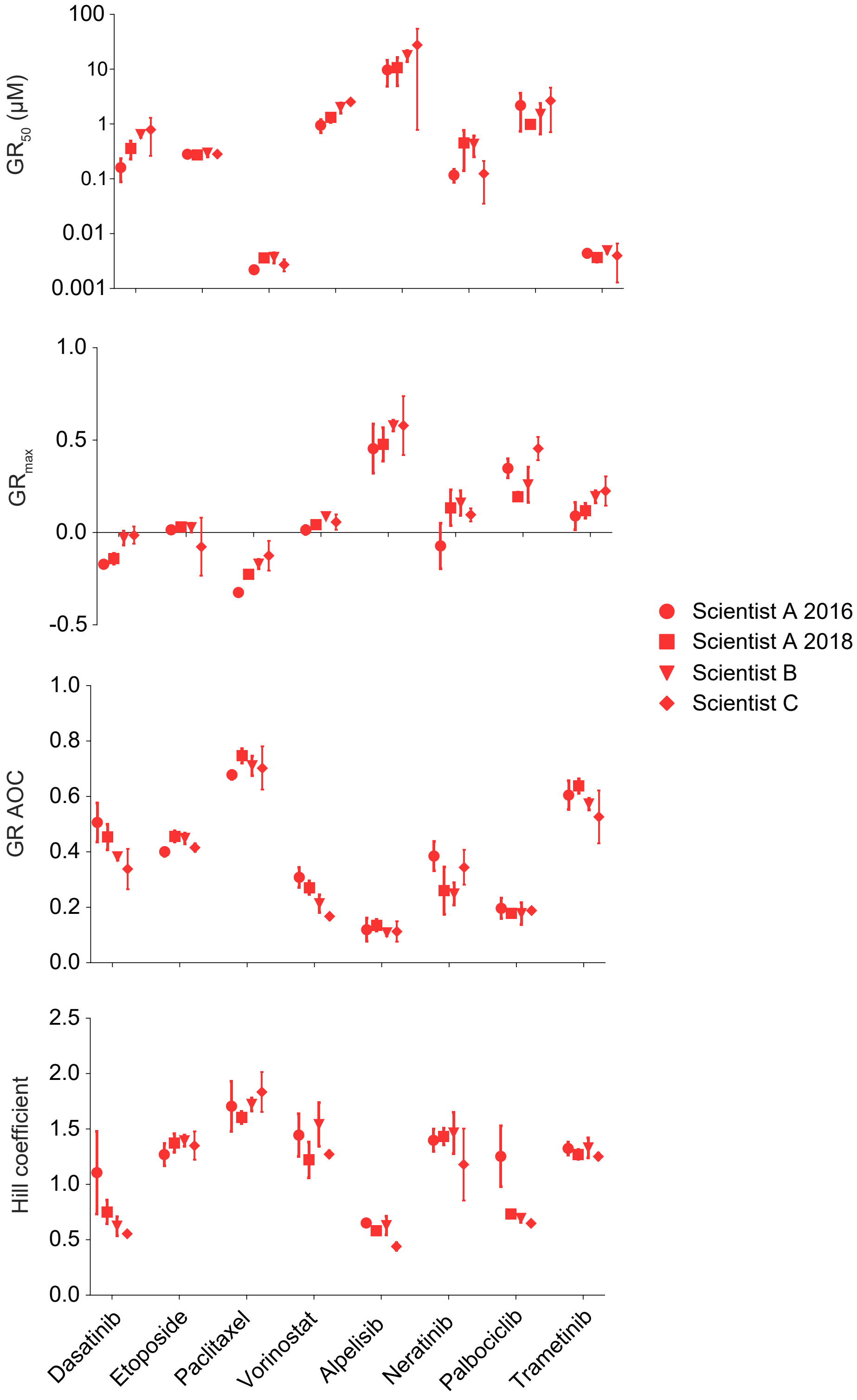

Figure S2, Related to Figure 2

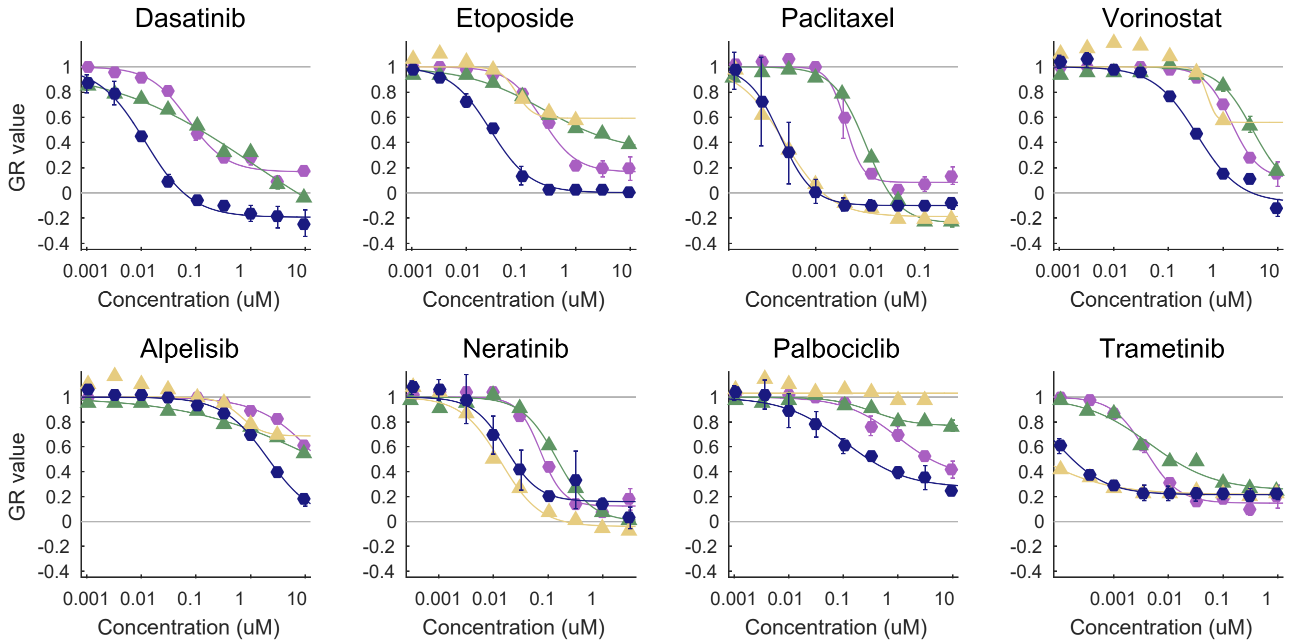

| | Drug dilution performed at: | Cell culture performed at: | Readout assay: |
|---|---|---|---|
| Preliminary 1 | Center 3 | Center 3 | Imaging |
| Preliminary 2 | Center 3 | Center 4 | CTG |
| Center 3 | Center 3 | Center 3 | Imaging |
| Center 4 | Center 4 | Center 4 | CTG |

Figure S3, Related to Figure 3

A

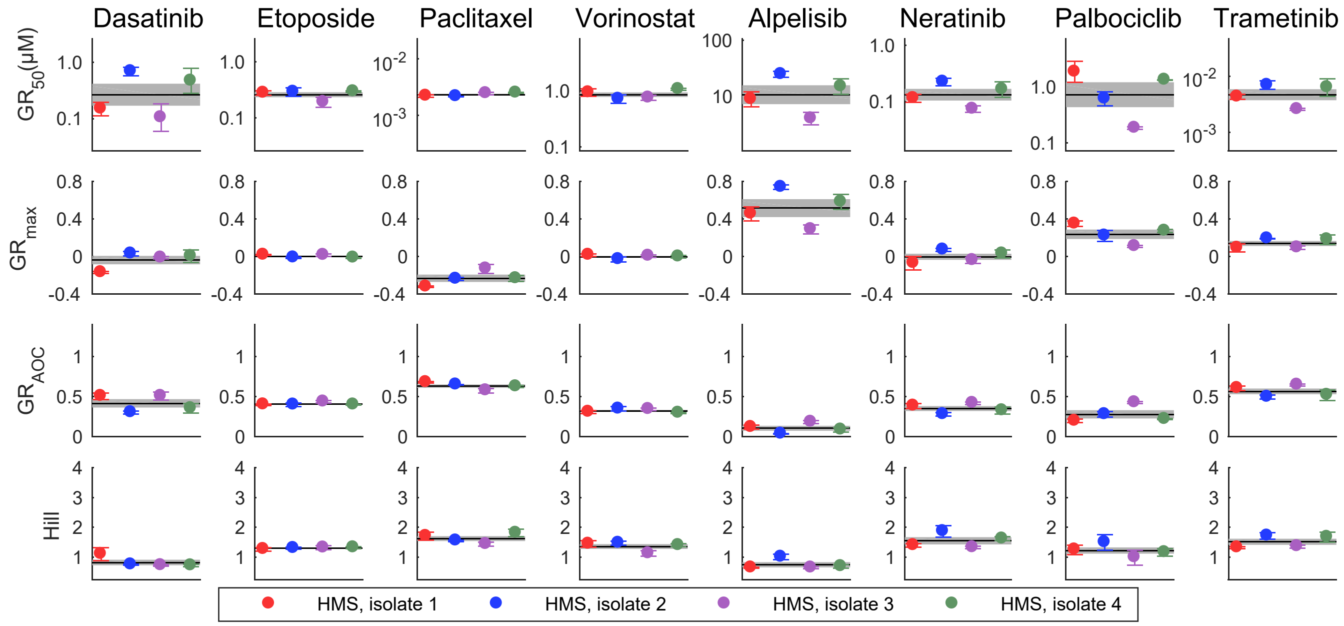

B

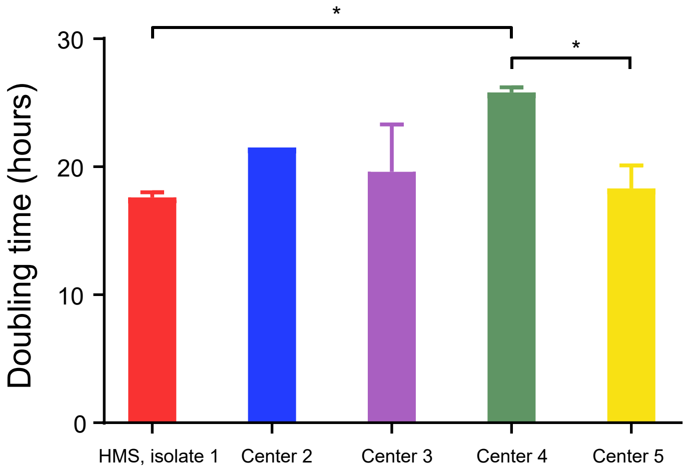

Figure S4, Related to Figure 3

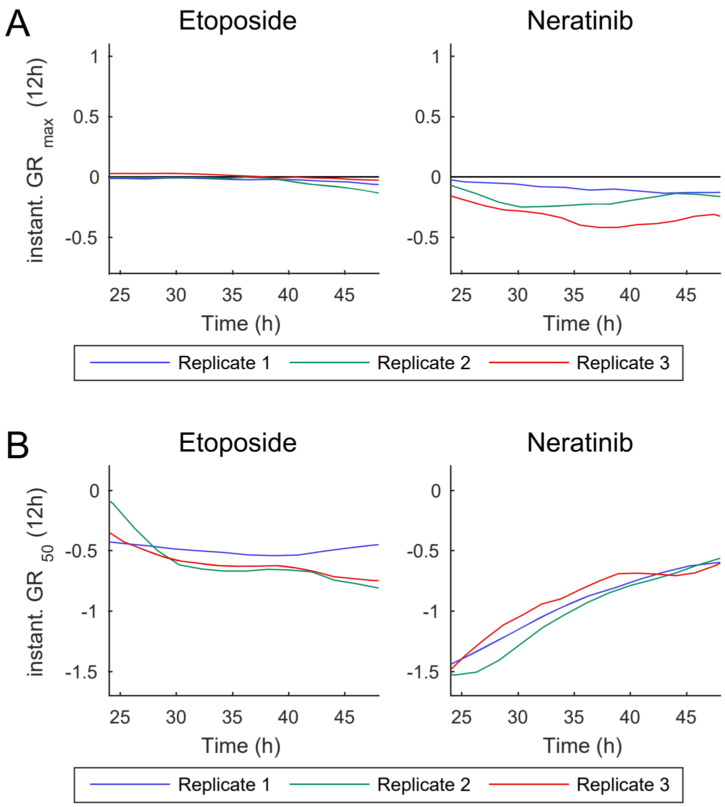

Figure S5, Related to Figure 4

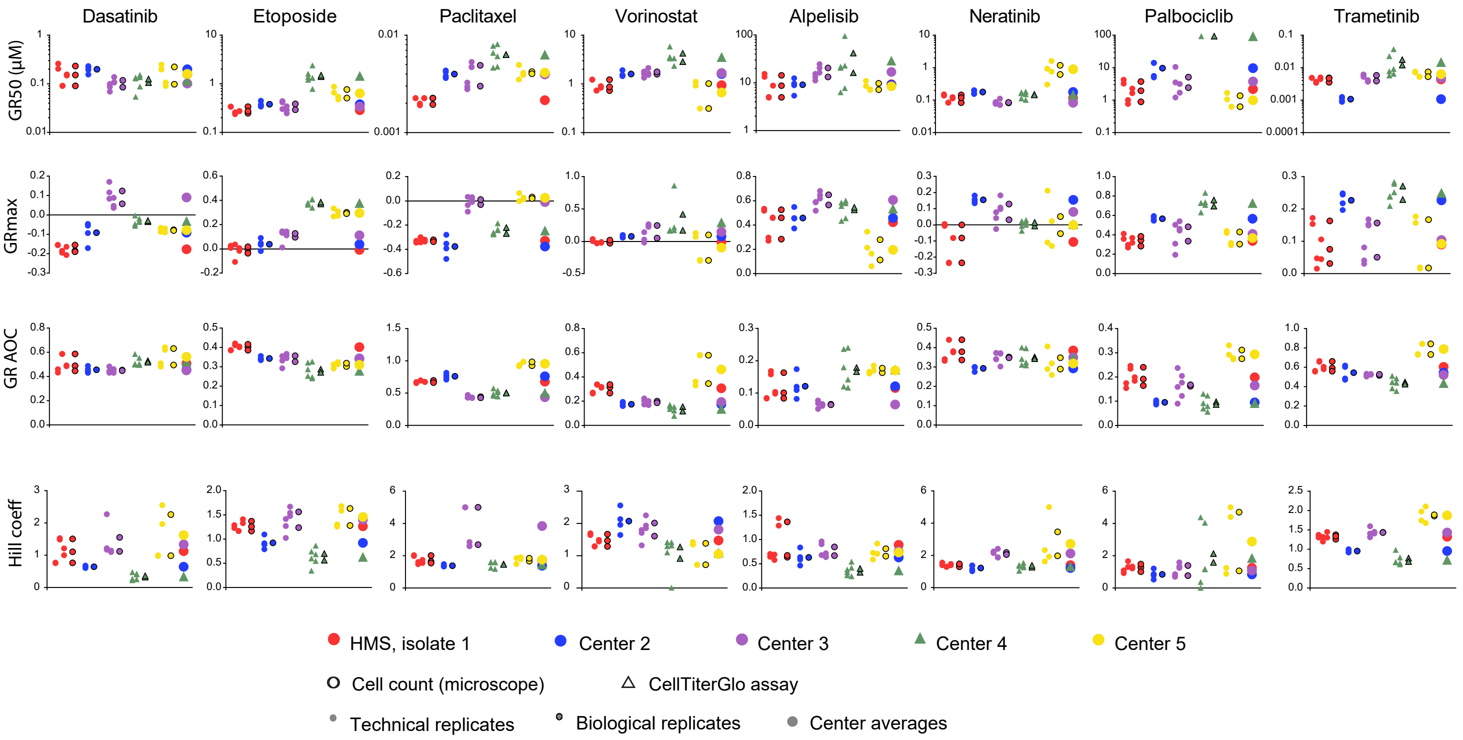

Figure S6, Related to Figure 6

Supplemental Table 1

### STR Profiling Results- GenePrint 10 Assay System- 3/31/16

| Sample Name | Comment | Marker | Allele 1 | Allele 2 | Allele 3 | Allele 4 | Allele 5 | Allele 6 | Peak Height Allele 1 | Peak Height Allele 2 | Peak Height Allele 3 | Peak Height Allele 4 | Peak Height Allele 5 |
|---|---|---|---|---|---|---|---|---|---|---|---|---|---|
| Allelic ladder | | TH01 | 4 | 5 | 6 | 7 | 8 | 9(more) | 634 | 521 | 593 | 516 | 591 |
| Allelic ladder | | D21S11 | 24 | 24.2 | 25 | 25.2 | 26 | 27(more) | 482 | 473 | 442 | 450 | 586 |
| Allelic ladder | | D5S818 | 7 | 8 | 9 | 10 | 11 | 12(more) | 897 | 641 | 582 | 760 | 667 |
| Allelic ladder | | D13S317 | 7 | 8 | 9 | 10 | 11 | 12(more) | 604 | 602 | 595 | 643 | 562 |
| Allelic ladder | | D7S820 | 6 | 7 | 8 | 9 | 10 | 11(more) | 746 | 720 | 638 | 639 | 631 |
| Allelic ladder | | D16S539 | 5 | 8 | 9 | 10 | 11 | 12(more) | 633 | 648 | 646 | 649 | 655 |
| Allelic ladder | | CSF1PO | 6 | 7 | 8 | 9 | 10 | 11(more) | 590 | 558 | 582 | 566 | 704 |
| Allelic ladder | | Amelogenin | X | Y | | | | | 557 | 629 | | | |
| Allelic ladder | | vWA | 10 | 11 | 12 | 13 | 14 | 15(more) | 489 | 582 | 599 | 493 | 514 |
| Allelic ladder | | TPOX | 6 | 7 | 8 | 9 | 10 | 11(more) | 507 | 594 | 570 | 543 | 544 |
| NTC | no template control | TH01 | 0 | 0 | | | | | | | | | |
| NTC | no template control | D21S11 | 0 | 0 | | | | | | | | | |
| NTC | no template control | D5S818 | 0 | 0 | | | | | | | | | |
| NTC | no template control | D13S317 | 0 | 0 | | | | | | | | | |
| NTC | no template control | D7S820 | 0 | 0 | | | | | | | | | |
| NTC | no template control | D16S539 | 0 | 0 | | | | | | | | | |
| NTC | no template control | CSF1PO | 0 | 0 | | | | | | | | | |
| NTC | no template control | Amelogenin | | | | | | | | | | | |
| NTC | no template control | vWA | 0 | 0 | | | | | | | | | |
| NTC | no template control | TPOX | 0 | 0 | | | | | | | | | |
| 2800M_pos_ctrl | positive control | TH01 | 6 | 9.3 | | | | | 2105 | 2151 | | | |
| 2800M_pos_ctrl | positive control | D21S11 | 29 | 31.2 | | | | | 1716 | 1751 | | | |
| 2800M_pos_ctrl | positive control | D5S818 | 12 | | | | | | 6411 | | | | |
| 2800M_pos_ctrl | positive control | D13S317 | 9 | 11 | | | | | 1919 | 1757 | | | |
| 2800M_pos_ctrl | positive control | D7S820 | 8 | 11 | | | | | 2650 | 2704 | | | |
| 2800M_pos_ctrl | positive control | D16S539 | 9 | 13 | | | | | 2911 | 2766 | | | |
| 2800M_pos_ctrl | positive control | CSF1PO | 12 | | | | | | 6539 | | | | |
| 2800M_pos_ctrl | positive control | Amelogenin | X | Y | | | | | 1692 | 1659 | | | |
| 2800M_pos_ctrl | positive control | vWA | 16 | 19 | | | | | 724 | 696 | | | |
| 2800M_pos_ctrl | positive control | TPOX | 11 | | | | | | 2550 | | | | |
| MCF10A-HMS_1 | replicate 1 | TH01 | 8 | 9.3 | | | | | 309 | 238 | | | |
| MCF10A-HMS_1 | replicate 1 | D21S11 | 28 | 30 | | | | | 295 | 259 | | | |
| MCF10A-HMS_1 | replicate 1 | D5S818 | 10 | 13 | | | | | 529 | 1018 | | | |
| MCF10A-HMS_1 | replicate 1 | D13S317 | 8 | 9 | | | | | 280 | 349 | | | |
| MCF10A-HMS_1 | replicate 1 | D7S820 | 10 | 11 | | | | | 609 | 525 | | | |
| MCF10A-HMS_1 | replicate 1 | D16S539 | 11 | 12 | | | | | 358 | 379 | | | |
| MCF10A-HMS_1 | replicate 1 | CSF1PO | 10 | 12 | | | | | 496 | 945 | | | |
| MCF10A-HMS_1 | replicate 1 | Amelogenin | X | | | | | | 488 | | | | |

### STR Profiling Results- GenePrint 10 Assay System- 3/31/16

| Sample Name | Comment | Marker | Allele 1 | Allele 2 | Allele 3 | Allele 4 | Allele 5 | Allele 6 | Peak Height Allele 1 | Peak Height Allele 2 | Peak Height Allele 3 | Peak Height Allele 4 | Peak Height Allele 5 |
|---|---|---|---|---|---|---|---|---|---|---|---|---|---|
| MCF10A-HMS_1 | replicate 1 | vWA | 15 | 17 | | | | | 132 | 118 | | | |
| MCF10A-HMS_1 | replicate 1 | TPOX | 9 | 11 | | | | | 191 | 155 | | | |
| MCF10A-HMS_2 | replicate 2 | TH01 | 8 | 9.3 | | | | | 294 | 317 | | | |
| MCF10A-HMS_2 | replicate 2 | D21S11 | 28 | 30 | | | | | 327 | 298 | | | |
| MCF10A-HMS_2 | replicate 2 | D5S818 | 10 | 13 | | | | | 550 | 1046 | | | |
| MCF10A-HMS_2 | replicate 2 | D13S317 | 8 | 9 | | | | | 347 | 314 | | | |
| MCF10A-HMS_2 | replicate 2 | D7S820 | 10 | 11 | | | | | 665 | 429 | | | |
| MCF10A-HMS_2 | replicate 2 | D16S539 | 11 | 12 | | | | | 532 | 560 | | | |
| MCF10A-HMS_2 | replicate 2 | CSF1PO | 10 | 12 | | | | | 568 | 1096 | | | |
| MCF10A-HMS_2 | replicate 2 | Amelogenin | X | | | | | | 489 | | | | |
| MCF10A-HMS_2 | replicate 2 | vWA | 15 | 17 | | | | | 173 | 112 | | | |
| MCF10A-HMS_2 | replicate 2 | TPOX | 9 | 11 | | | | | 152 | 192 | | | |
| MCF10A-H2B-mCherry_1 | replicate 1 | TH01 | 8 | 9.3 | | | | | 288 | 229 | | | |
| MCF10A-H2B-mCherry_1 | replicate 1 | D21S11 | 28 | 30 | | | | | 272 | 276 | | | |
| MCF10A-H2B-mCherry_1 | replicate 1 | D5S818 | 10 | 13 | | | | | 363 | 904 | | | |
| MCF10A-H2B-mCherry_1 | replicate 1 | D13S317 | 8 | 9 | | | | | 205 | 273 | | | |
| MCF10A-H2B-mCherry_1 | replicate 1 | D7S820 | 10 | 11 | | | | | 341 | 420 | | | |
| MCF10A-H2B-mCherry_1 | replicate 1 | D16S539 | 11 | 12 | | | | | 400 | 371 | | | |
| MCF10A-H2B-mCherry_1 | replicate 1 | CSF1PO | 10 | 12 | | | | | 520 | 807 | | | |
| MCF10A-H2B-mCherry_1 | replicate 1 | Amelogenin | X | | | | | | 353 | | | | |
| MCF10A-H2B-mCherry_1 | replicate 1 | vWA | 15 | 17 | | | | | 104 | 97 | | | |
| MCF10A-H2B-mCherry_1 | replicate 1 | TPOX | 9 | 11 | | | | | 140 | 122 | | | |
| MCF10A-H2B-mCherry_2 | replicate 2 | TH01 | 8 | 9.3 | | | | | 178 | 183 | | | |
| MCF10A-H2B-mCherry_2 | replicate 2 | D21S11 | 28 | 30 | | | | | 200 | 197 | | | |
| MCF10A-H2B-mCherry_2 | replicate 2 | D5S818 | 10 | 13 | | | | | 256 | 495 | | | |
| MCF10A-H2B-mCherry_2 | replicate 2 | D13S317 | 8 | 9 | | | | | 158 | 227 | | | |
| MCF10A-H2B-mCherry_2 | replicate 2 | D7S820 | 10 | 11 | | | | | 380 | 374 | | | |
| MCF10A-H2B-mCherry_2 | replicate 2 | D16S539 | 11 | 12 | | | | | 262 | 265 | | | |
| MCF10A-H2B-mCherry_2 | replicate 2 | CSF1PO | 10 | 12 | | | | | 283 | 664 | | | |
| MCF10A-H2B-mCherry_2 | replicate 2 | Amelogenin | X | | | | | | 230 | | | | |
| MCF10A-H2B-mCherry_2 | replicate 2 | vWA | 15 | 17 | | | | | 79 | 76 | | | |
| MCF10A-H2B-mCherry_2 | replicate 2 | TPOX | 9 | 11 | | | | | 125 | 94 | | | |
| MCF10A-OHSU_1 | replicate 1 | TH01 | 8 | 9.3 | | | | | 195 | 236 | | | |
| MCF10A-OHSU_1 | replicate 1 | D21S11 | 28 | 30 | | | | | 236 | 217 | | | |
| MCF10A-OHSU_1 | replicate 1 | D5S818 | 10 | 13 | | | | | 362 | 766 | | | |
| MCF10A-OHSU_1 | replicate 1 | D13S317 | 8 | 9 | | | | | 220 | 257 | | | |
| MCF10A-OHSU_1 | replicate 1 | D7S820 | 10 | 11 | | | | | 430 | 433 | | | |
| MCF10A-OHSU_1 | replicate 1 | D16S539 | 11 | 12 | | | | | 399 | 262 | | | |
| MCF10A-OHSU_1 | replicate 1 | CSF1PO | 10 | 12 | | | | | 428 | 840 | | | |

### STR Profiling Results- GenePrint 10 Assay System- 3/31/16

| Sample Name | Comment | Marker | Allele 1 | Allele 2 | Allele 3 | Allele 4 | Allele 5 | Allele 6 | Peak Height Allele 1 | Peak Height Allele 2 | Peak Height Allele 3 | Peak Height Allele 4 | Peak Height Allele 5 |
|---|---|---|---|---|---|---|---|---|---|---|---|---|---|
| MCF10A-OHSU_1 | replicate 1 | Amelogenin | X | | | | | | 397 | | | | |
| MCF10A-OHSU_1 | replicate 1 | vWA | 15 | 17 | | | | | 88 | 111 | | | |
| MCF10A-OHSU_1 | replicate 1 | TPOX | 9 | 11 | | | | | 130 | 125 | | | |
| MCF10A-OHSU_2 | replicate 2 | TH01 | 8 | 9.3 | | | | | 267 | 261 | | | |
| MCF10A-OHSU_2 | replicate 2 | D21S11 | 28 | 30 | | | | | 291 | 313 | | | |
| MCF10A-OHSU_2 | replicate 2 | D5S818 | 10 | 13 | | | | | 500 | 1000 | | | |
| MCF10A-OHSU_2 | replicate 2 | D13S317 | 8 | 9 | | | | | 262 | 328 | | | |
| MCF10A-OHSU_2 | replicate 2 | D7S820 | 10 | 11 | | | | | 622 | 407 | | | |
| MCF10A-OHSU_2 | replicate 2 | D16S539 | 11 | 12 | | | | | 396 | 368 | | | |
| MCF10A-OHSU_2 | replicate 2 | CSF1PO | 10 | 12 | | | | | 431 | 789 | | | |
| MCF10A-OHSU_2 | replicate 2 | Amelogenin | X | | | | | | 485 | | | | |
| MCF10A-OHSU_2 | replicate 2 | vWA | 15 | 17 | | | | | 157 | 150 | | | |
| MCF10A-OHSU_2 | replicate 2 | TPOX | 9 | 11 | | | | | 231 | 201 | | | |
| MCF10A-GM_1 | replicate 1 | TH01 | 8 | 9.3 | | | | | 256 | 196 | | | |
| MCF10A-GM_1 | replicate 1 | D21S11 | 28 | 30 | | | | | 297 | 252 | | | |
| MCF10A-GM_1 | replicate 1 | D5S818 | 10 | 13 | | | | | 628 | 1394 | | | |
| MCF10A-GM_1 | replicate 1 | D13S317 | 8 | 9 | | | | | 431 | 366 | | | |
| MCF10A-GM_1 | replicate 1 | D7S820 | 10 | 11 | | | | | 679 | 463 | | | |
| MCF10A-GM_1 | replicate 1 | D16S539 | 11 | 12 | | | | | 519 | 497 | | | |
| MCF10A-GM_1 | replicate 1 | CSF1PO | 10 | 12 | | | | | 654 | 1266 | | | |
| MCF10A-GM_1 | replicate 1 | Amelogenin | X | | | | | | 729 | | | | |
| MCF10A-GM_1 | replicate 1 | vWA | 15 | 17 | | | | | 149 | 159 | | | |
| MCF10A-GM_1 | replicate 1 | TPOX | 9 | 11 | | | | | 174 | 177 | | | |
| MCF10A-GM_2 | replicate 2 | TH01 | 8 | 9.3 | | | | | 452 | 389 | | | |
| MCF10A-GM_2 | replicate 2 | D21S11 | 28 | 30 | | | | | 505 | 576 | | | |
| MCF10A-GM_2 | replicate 2 | D5S818 | 10 | 13 | | | | | 943 | 1395 | | | |
| MCF10A-GM_2 | replicate 2 | D13S317 | 8 | 9 | | | | | 489 | 461 | | | |
| MCF10A-GM_2 | replicate 2 | D7S820 | 10 | 11 | | | | | 890 | 755 | | | |
| MCF10A-GM_2 | replicate 2 | D16S539 | 11 | 12 | | | | | 713 | 591 | | | |
| MCF10A-GM_2 | replicate 2 | CSF1PO | 10 | 12 | | | | | 676 | 1415 | | | |
| MCF10A-GM_2 | replicate 2 | Amelogenin | X | | | | | | 893 | | | | |
| MCF10A-GM_2 | replicate 2 | vWA | 15 | 17 | | | | | 268 | 210 | | | |
| MCF10A-GM_2 | replicate 2 | TPOX | 9 | 11 | | | | | 240 | 278 | | | |

#### STR Profiling Results- GenePrint 10 Assay System- 3/31/16

| Peak Height Allele 6 |
| --- |
| 533 |
| 498 |
| 759 |
| 644 |
| 579 |
| 645 |
| 739 |
| 513 |
| 507 |

### STR Profiling Results- GenePrint 10 Assay System- 3/31/16

| Peak Height Allele 6 |
|---|

### STR Profiling Results- GenePrint 10 Assay System- 3/31/16

| Peak Height Allele 6 |
| --- |

Supplemental Table 2

| **Reagent** | **Source** | **Catalogue number** | **Lot number** |
|---|---|---|---|
| *Media components* | | | |
| DMEM/F12 | Invitrogen | 11330-032 | 1747275 |
| Horse serum | Sigma | H1138 | 12B496 |
| rhEGF | R&D Systems | 236-EG | HLM7515071 |
| Hydrocortisone | Sigma | H4001 | SLBN5690V |
| Cholera toxin | Sigma | C8052 | 095M4093V |
| Insulin | Sigma | I9278 | SLBP1369V |
| Pen/Strep | Invitrogen | 15070-063 | 1697552 |
| *Small molecules* | | | |
| Alpelisib | MedChem Express | HY-15244 | 06192 |
| Dasatinib | MedChem Express | HY-10191 | 13044 |
| Etoposide | MedChem Express | HY-13629 | 11793 |
| Neratinib | MedChem Express | HY-32721 | 10283 |
| Paclitaxel | MedChem Express | HY-B0015 | 18138 |
| Palbociclib | MedChem Express | HY-50767 | 16349 |
| Trametinib | MedChem Express | HY-10999 | 07378 |
| Vorinostat | MedChem Express | HY-10221 | 09386 |
